# Songs distinguish the cryptic giant hummingbird species and clarify range limits

**DOI:** 10.1101/2025.06.30.662449

**Authors:** Bryce W. Robinson, Ryan J. Zucker, Christopher C. Witt, Thomas Valqui, Jessie L. Williamson

## Abstract

Vocal traits are often essential for distinguishing phenotypically cryptic taxa. The hummingbird genus *Patagona* comprises two species, near identical in plumage and morphology, that differ in almost every aspect of their ecology and evolution: The Northern Giant Hummingbird (*Patagona peruviana*) and Southern Giant Hummingbird (*Patagona gigas*). Here, we characterized the songs of both giant hummingbird species and assessed whether song can be used to distinguish the two in the field. We recorded both species in Peru, Bolivia, and Chile in 2023 and 2025 and used public data to analyze song variation of 217 individuals recorded across the Andes. Sampling spanned 49 years, >36° of latitude, and >4,300 meters in elevation. We first quantified species-level song differences in allopatric breeding populations and trained a linear discriminant model to identify individuals to species. The trained model had 100% classification accuracy. We then used our trained model to identify individuals recorded during co-occurring, non-breeding periods of overlap, and subsequently analyzed range-wide song variation; this model had 98.72% classification accuracy (1.28% error rate; one individual misidentified). We found striking song divergence between the two species, uncovering that Northern and Southern Giant Hummingbirds can be reliably and easily identified by song across their ranges and during any month of the year. We provide new data on the range limits of both species in northern Bolivia, highlighting a previously unknown zone of overlap around Lake Titicaca. Unlike other phenotypic traits, song provides a robust method for identifying the giant hummingbird species, opening doors to future research on ecology, trait evolution, hybrid zone dynamics, and conservation of the world’s largest hummingbirds.

## INTRODUCTION

Numerous bird species differ only subtly in external phenotype but produce strikingly different sounds. These acoustic traits, including song, are principal signals used in mate choice and species recognition, making vocal traits central to understanding reproductive isolation and speciation (Catchpole and Slater 1995). Acoustic traits are also essential for field identification: For example, previous work has shown that vocalizations can distinguish cryptic species of *Rallus* rails (Stiffler et al. 2018), *Glaucidium* owlets (Gwee et al. 2019), *Bubo* owls (Movin et al. 2022), *Tanysiptera* kingfishers (Sin et al. 2022), *Sclerurus* leaftossers (Cooper and Cuervo 2017), *Troglodytes* wrens (Toews and Irwin 2008), *Camptostoma* tyrannulets (Lima and Vaz 2024), and *Phylloscopus* warblers (Irwin 2000), as well as species of fish (Parmentier et al. 2022), frogs (Wilczynski et al. 1993), and nocturnal primates (Braune et al. 2008). Characterizing vocal traits therefore provides a powerful tool for identifying cryptic taxa, guiding research on ecology, taxonomy, and evolutionary divergence (Toews and Irwin 2008; Ng and Rheindt 2016; Stiffler et al. 2018; Gwee et al. 2019; Movin et al. 2022; Sin et al. 2022).

The hummingbirds (family Trochilidae) are one of three bird families that independently evolved vocal learning (Jarvis et al. 2000; Hackett et al. 2008). Yet, hummingbirds are rarely studied for their songs, perhaps because most species produce largely single-note songs (e.g., (Atwood et al. 1991; Araya-Salas and Wright 2013). However, a number of hummingbird species have intermediate vocal complexity (Baptista and Schuchmann 1990; Gaunt et al. 1994) and some rival passerines in their phonological and syntactic song intricacy (e.g., Ficken et al. 2000; Jarvis et al. 2000; Ornelas et al. 2002; González and Ornelas 2005, 2009; Ferreira et al. 2006; González et al. 2011). The diversity of vocal repertoires, independent evolution of song learning, and ability to learn complex songs makes the hummingbirds an interesting family through which to study song differences in sister taxa.

The giant hummingbird clade (genus *Patagona*) contains two species that diverged 2.1–3.4 Mya and that differ in breeding latitude, migratory behavior, and respiratory physiology (Williamson and Witt 2021a; Williamson et al. 2023, 2024). While the two differ in average plumage patterns and external measurements, these differences are insufficient to identify all individuals (Williamson et al. 2025). Both giant hummingbird species are found in arid and semi-arid habitats and in intermontane valleys of the Andes, including scrub, hedgerows, agricultural areas, open woods (including *Polylepis*), and gardens, often with *Agave*, *Puya*, and Cactaceae spp. Across their ranges, they tend to associate with fast-flowing streams. The Northern Giant Hummingbird (*Patagona peruviana*) is a high-elevation resident known to breed in Ecuador, Peru, and northern Chile that occurs at ∼1,800–4,300 meters (m) throughout the north-central Andes (Ridgely and Greenfield 2001; Jaramillo 2003; Schulenberg et al. 2010; Williamson et al. 2025). The Southern Giant Hummingbird (*Patagona gigas*) breeds from sea level to ∼2,500 m in central Chile, above ∼2,500 m in northwest Argentina, and ∼2,100-4,100 m in Bolivia. Southern latitude populations are long-distance migrants: During the austral winter (∼March–Sept), Chilean-breeding Southern Giant Hummingbirds migrate north to tropical latitudes of the central Peruvian Andes where individuals overlap extensively with populations of the Northern Giant Hummingbird (Williamson et al. 2024). At least some breeding populations from northwest Argentina also migrate to central Peru, but further details about migration are currently unknown. The two species seasonally co-occur at the same sites and elevations during the nonbreeding season.

While the giant hummingbirds have diverged in many aspects of their ecology and evolution, they are phenotypically cryptic and difficult to distinguish––a challenge that has hindered attempts to characterize their range limits and seasonal co-occurrence (Williamson et al. 2024, 2025). The two species have near-identical body mass distributions and differ only subtly in plumage, with a few key features providing evidence for species identity: Throat color and pattern, eye-ring, post-ocular spot, wing length, bill length, and tail length tend to be informative (Williamson et al. 2025). However, even among adults, the range of plumage variation, particularly in throat color and pattern, overlaps within and between species (Williamson et al. 2024). While the Northern Giant Hummingbird averages larger in bill length, wing chord, tail length, and tarsus length, there is substantial overlap with the Southern Giant Hummingbird, such that only ∼65–85% of individuals can be identified to species using measurements when comparing sexes separately and with a trained linear discriminant model (Williamson et al. 2024, 2025). Many individuals are thus best left unidentified when observed at sites and during seasons of co-occurrence.

Although the two species are largely allopatric during their respective breeding seasons, Williamson et al. (2024) reported a male Northern x Southern Giant Hummingbird F1 hybrid from Lima Department, Peru, in September. The hybrid had Peruvian and Argentinian ancestry, and its occurrence suggests breeding range overlap between the Northern and Southern Giant Hummingbird (Williamson et al. 2024). These findings raise questions about seasonal range limits and the potential roles of song in species recognition and field identification.

Both giant hummingbird species sing simple, single-element songs, often referred to as ‘calls’, which have been previously described as “a high-pitched *cwueet*” (Ridgely and Greenfield 2001), “a squeaky, loud *heeee*” (Schulenberg et al. 2010), a “characteristic melancholy pipping note” (Koepcke 1983), “a squeaky *zeet*” (Herzog et al. 2016), “a piercing *SIP* whistle” (Pearman and Areta 2020), and “a loud single chip note” (Jaramillo 2003). The two make a variety of other vocalizations and chatter calls during both the breeding and non-breeding seasons, described as “squeaky whistles and trills” (Velásquez-Noriega et al. 2023). Both sing from exposed perches, as well as in flight and during aggressive in-flight chases with congeners. Although previous work has suggested that some species of hummingbirds may be open-ended song learners, capable of modifying songs into adulthood (Araya-Salas and Wright 2013; Johnson and Clark 2020), neither song learning capability nor vocal repertoire has been studied in the giant hummingbirds, and *Patagona* remains the only major hummingbird clade for which no systematic characterization of vocalizations has ever been conducted (Duque and Carruth 2022).

Here, we used range-wide comparative analysis of vocal traits to characterize the songs of the two giant hummingbird species, and we evaluated whether song can be used to distinguish the two––particularly in non-breeding zones of overlap. We collected field recordings of singing giant hummingbirds during both the breeding and nonbreeding season from Peru, Bolivia, and Chile, which we combined with recordings from public repositories to study song trait variation across the range of the genus. We predicted finding subtle variation in both species’ single-element songs that mirrors the level of divergence observed in phenotypic comparisons. Accordingly, we posited that models trained to classify songs to species-level would have similar low to moderate predictive power as models previously trained using plumage and morphology (Williamson et al. 2024).

## METHODS

### Field recording

In 2023 and 2025, JLW collected 53 recordings of the two giant hummingbird species during both opportunistic and targeted fieldwork in Peru, Chile, and Bolivia. Fieldwork took place in Lima and Ancash Departments of Peru in July–Aug 2023, Lima Department in Jan 2025, in the Valparaíso Region of central Chile during Jan–Feb 2025, and in La Paz and Cochabamba Departments of Bolivia during Feb 2025. During field recording, JLW visited sites where giant hummingbirds had been recently seen or reported (per the eBird database and suggestions from local guides), as well as sites whose habitat appeared suitable for one or both species. Upon finding a focal giant hummingbird, JLW carefully observed the individual and took photographs and recordings, to the extent possible. Most recordings (>81%) were captured with a Zoom F3 Field Recorder and Sennheiser ME67 shotgun microphone, but several were taken using an iPhone 14 Pro with either the Voice Memos or Merlin apps. All recordings, as well as representative photographs, were later archived in the Macaulay Library (ML; https://www.macaulaylibrary.org/) at the Cornell Lab of Ornithology.

### Song data collection

We compiled a total of 217 song recordings of Northern and Southern Giant Hummingbirds that were made between 1976–2025, during months that spanned the breeding and non-breeding seasons of both species. Of these, 189 recordings were from the Macaulay Library and 28 from Xeno-canto (XC; https://www.xeno-canto.org). All data available as of 15 February 2025 were used. Once aggregated, we compared recordist names, recording dates, and localities of Macaulay Library versus Xeno-canto data to ensure each recording was unique. We georeferenced missing latitudes and longitudes for 14 recordings missing coordinates, as described in Williamson and Witt 2021b. Each recording contained a principal, single-element song (Fig. 1); many also contained a variety of flight and chatter calls. Because calls varied substantially in length, pitch, and frequency and were often given in response to interactions with conspecifics or other species, we analyzed only single-element songs.

**Figure 1.**
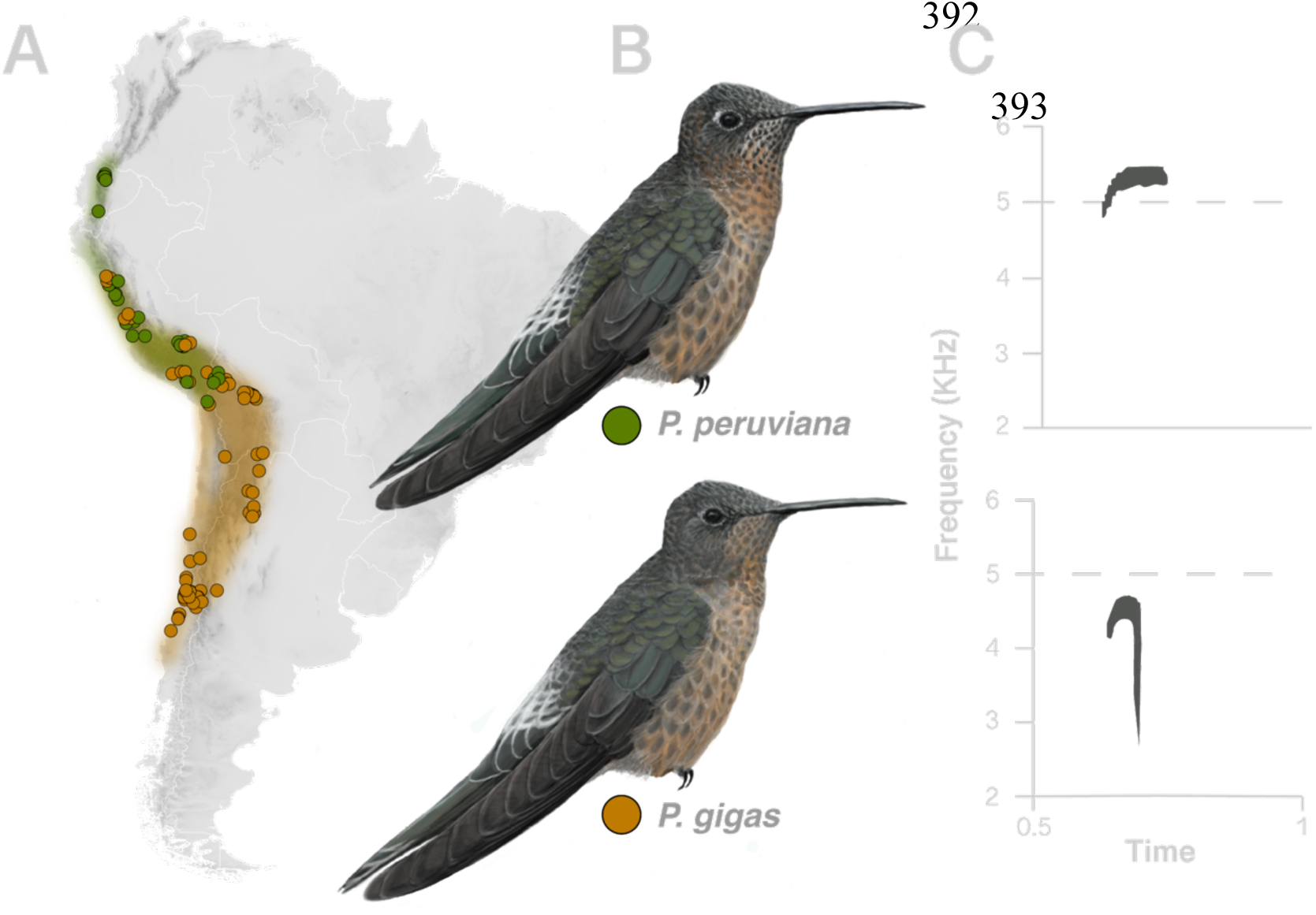
Recordings from across the Andes were analyzed to characterize the songs of the two cryptic giant hummingbird species. **A** Estimated annual ranges and non-breeding seasonal overlap of the Northern Giant Hummingbird (*Patagona peruviana*) and Southern Giant Hummingbird (*Patagona gigas*). Points indicate the localities of 217 analyzed recordings spanning 1976–2025, obtained from the Macaulay Library at the Cornell Lab of Ornithology (https://www.macaulaylibrary.org/) and XenoCanto). Points were colored by a priori analysis of season, locality, and date, and later confirmed with linear discriminant analysis classification. **B** The two giant hummingbird species differ only subtly in plumage. Illustrations: Bryce W. Robinson. **C** Spectrograms illustrate vocal differences (top=*P. peruviana*; bottom: *P. gigas*).

### Song processing and annotation

We measured the following variables in spectrograms from .wav, .mp3, and .m4a files of each song recording: Start time (secs), end time (secs), minimum or lowest frequency in kilohertz (Khz), and maximum or highest frequency (Khz). We scored three songs (i.e., unique elements) per recording of each individual, and when possible, scored three consecutive songs. When fewer than three songs were available or scorable, we analyzed all scorable songs. We assigned each a coarse quality score (“low”, “medium”, “high”) and noted whether audio files were compressed or noise reduced. We excluded recordings from our analysis that contained disruptive background noise, spectrographic artifacts, strong echoes, and atypical chatter notes (52 recordings total, or ∼24%). To ensure consistency, RJZ conducted all scoring in Raven Pro v.1.6.5 (K. Lisa Yang Center for Conservation Bioacoustics at the Cornell Lab of Ornithology 2024).

Using raw data scores for each recording, we calculated song duration (measured as song end time minus song start time; in secs) and frequency range, sometimes called ‘bandwidth’ (measured as maximum frequency minus minimum frequency), two commonly used parameters in vocal analysis (Odom et al. 2021). For each individual, we then calculated a mean from the measurements of all scored songs for each parameter, including: 1) minimum frequency, 2) maximum frequency, 3) song duration, and 4) frequency bandwidth, as described in (Toews and Irwin 2008). For 27 individuals, it was necessary to calculate mean trait values from only two scored songs; and for eight individuals, we took scores from a single song. We then used mean measurements for each individual in downstream analyses.

We divided the data into breeding and full annual (including the breeding and non-breeding seasons) subsets for separate analyses. Breeding data enabled us to characterize the songs of the giant hummingbirds using individuals of known species identity, while full annual data enabled us to test how our model performed when identifying individuals of unknown species identity. In the breeding dataset, species identity was determined using a combination of month, locality, and if possible, plumage characteristics from accompanying photos taken in the field or available in eBird checklists. We analyzed the distributions of expected song traits within each the Northern and Southern Giant Hummingbird and dropped seven outlier individuals (four Northerns and three Southerns) that fell outside the expected trait distributions for each species. For the Northern Giant Hummingbird, these included: Recordings from two Northerns that had longer-than-expected song durations (ML242924 and ML33867; individual ML33867 had only a single scored song); and two individuals with exceptionally low minimum frequency, low maximum frequency, short song duration, and low frequency range (the first, ML511071761, a putative late migrant Southern Giant Hummingbird, had only one scored song and was recorded 25 October 2022 from Cusco, Peru; the second, XC17922, a putative early migrant Southern Giant Hummingbird, was recorded 9 March 2006 from Cusco, Peru). Southern Giant Hummingbird outliers included: A recording from one individual with a low maximum song frequency (ML625624959), and two individuals with longer-than-expected song durations (ML410550051 and ML69741521; the latter was also classified as having low song quality). We conservatively excluded recordings from Bolivia from our breeding range dataset due to uncertainty about the distributions of giant hummingbird species in the central Andes (Williamson et al. 2025). Our final breeding range dataset included songs from 25 Northern and 55 Southern Giant Hummingbirds.

### Song analysis

We first visually examined song characteristics of both the Northern and Southern Giant Hummingbird using spectrograms, noting differences in shape, pacing, and frequency. We used *t*-tests or non-parametric Wilcoxon signed-rank tests to quantitatively compare differences in mean song characteristics (minimum frequency, maximum frequency, duration, and frequency range) between the two species. Next, to test the reliability of species recognition by song, we standardized mean song traits and ran a Principal Components Analysis (PCA) with means of minimum song frequency, maximum song frequency, and frequency range. We excluded song duration from PCA because of a high correlation with both mean minimum and maximum song frequency, the metrics from which mean duration was calculated. With these same three standardized traits, we then conducted a linear discriminant analysis (LDA), using a randomized 80% of breeding season data for model training and 20% for model testing, without replacement. All statistical tests and models were carried out using R (R Core Team 2019).

Our analyses confirmed strong differences between the songs of Northern and Southern Giant Hummingbirds in all measured characteristics and the trained LDA model was able to distinguish the two species with 100% accuracy using breeding data. We therefore used our trained LDA model, checked against qualitative assessment of spectrogram shapes, to classify each non-breeding season recording as either a Northern or Southern Giant Hummingbird. We then combined breeding and non-breeding season data from across the annual ranges of both species. After outlier assessment, our combined dataset included a total of 157 individuals (41 Northern Giant Hummingbirds and 116 Southern Giant Hummingbirds). We repeated comparative analyses of trait differences, as well as PCA and LDA analyses.

We additionally conducted LDA analyses using published morphological data (lengths of bill, wing, and tail) from Williamson et al. (2025) to compare the utility of song versus external phenotypic characters in distinguishing the two species. Our morphology dataset included 166 adult females and 161 adult males from across the annual range.

## RESULTS

### Comparative analysis of song traits

The songs of the Northern and Southern Giant Hummingbird differed significantly in all measured traits (Fig. 2; Table 1). Northern Giant Hummingbirds had significantly higher minimum (*p*=2.2×10^-16^, Wilcoxon rank sum test) and maximum (*p*=<2.2×10^-16^, *t*-test) song frequencies than Southern Giant Hummingbirds. The song of the Northern Giant Hummingbird was also 2x longer (*p*=2.2×10^-16^, Wilcoxon rank sum test). The song of the Southern Giant Hummingbird spanned a 2.4x greater frequency range than the song of the Northern Giant Hummingbird (*p*=2.2×10^-16^, Wilcoxon rank sum test). These differences were consistent when comparing breeding season-only and full annual data (Fig. 2).

**Figure 2.**
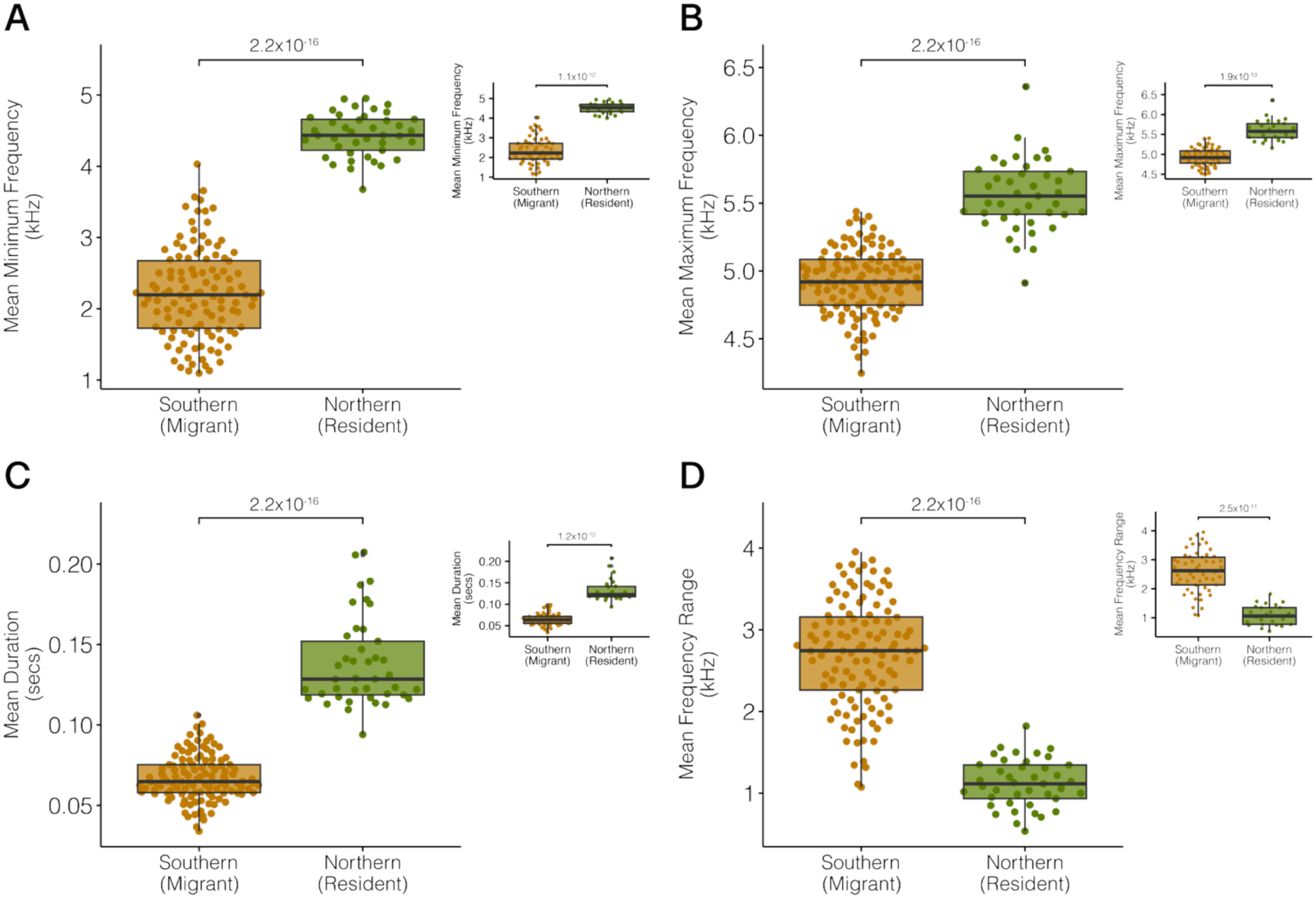
The songs of Northern and Southern Giant Hummingbirds differ in all measured characteristics. **A–D** Comparisons of song characteristics from full annual data (main plots; *n*=117 Southern songs, *n*=41 Northern songs) and breeding season-only data (insets; *n*=55 Southern songs, *n*=25 Northern songs). **A** Minimum frequency was 2x higher in Northern than Southern Giant Hummingbirds (*p* < 2.2×10^-16^, Wilcoxon rank-sum test). **B** Maximum frequency was 1.3x higher in Northern Giant Hummingbirds (*p* < 2.2×10^-16^, *t*-test). **C** Northern Giant Hummingbird songs are 2x longer than Southern Giant Hummingbird songs (*p* < 2.2×10^-16^, Wilcoxon rank-sum test). **D** Frequency bandwidth (range) was 2.41x greater in Southern Giant hummingbirds (*p* < 2.2×10^-16^, Wilcoxon rank-sum test). In all panels, box plot horizontal lines indicate median values. Each point is a value from a single song.

**Table 1.** Song trait summary statistics for each Northern and Southern Giant Hummingbirds (sexes undetermined). For each parameter, we provide sample size (*n*), mean, standard deviation, range (minimum and maximum values), and coefficient of variation.

| Species | Dataset | Parameter (units) | $n$ | Mean (kHz) | Standard Deviation | Minimum (kHz) | Maximum (kHz) | Coefficient of Variation |
| --- | --- | --- | --- | --- | --- | --- | --- | --- |
| Northern | Breeding | Mean min frequency | 25 | 4.52 | 0.28 | 4.01 | 4.95 | 0.06 |
|  |  | Mean max frequency | 25 | 5.6 | 0.27 | 5.16 | 6.36 | 0.05 |
|  |  | Mean duration | 25 | 0.13 | 0.03 | 0.09 | 0.21 | 0.2 |
|  |  | Mean frequency range | 25 | 1.09 | 0.34 | 0.53 | 1.82 | 0.31 |
|  | Annual | Mean min frequency | 41 | 4.44 | 0.3 | 3.68 | 4.95 | 0.07 |
|  |  | Mean max frequency | 41 | 5.56 | 0.26 | 4.91 | 6.36 | 0.05 |
|  |  | Mean duration | 41 | 0.14 | 0.03 | 0.09 | 0.21 | 0.2 |
|  |  | Mean frequency range | 41 | 1.12 | 0.29 | 0.53 | 1.82 | 0.26 |
| Southern | Breeding | Mean min frequency | 55 | 2.34 | 0.66 | 1.13 | 4.03 | 0.28 |
|  |  | Mean max frequency | 55 | 4.93 | 0.22 | 4.49 | 5.4 | 0.04 |
|  |  | Mean duration | 55 | 0.06 | 0.01 | 0.03 | 0.1 | 0.21 |
|  |  | Mean frequency range | 55 | 2.59 | 0.71 | 1.07 | 3.96 | 0.27 |
|  | Annual | Mean min frequency | 116 | 2.22 | 0.63 | 1.09 | 4.03 | 0.29 |
|  |  | Mean max frequency | 116 | 4.92 | 0.24 | 4.25 | 5.44 | 0.05 |
|  |  | Mean duration | 116 | 0.07 | 0.01 | 0.03 | 0.11 | 0.21 |
|  |  | Mean frequency range | 116 | 2.7 | 0.66 | 1.07 | 3.96 | 0.24 |

Quantitative song differences between species were consistent with visual inspections of the spectrograms (Fig. 1) and were easily audible when listening to recordings. In all scored spectrograms, the song of the Northern Giant Hummingbird appeared as an initial upward-sloping note leading to a short, horizontal segment; by ear, the song was recognizable as a high-pitched, dry and thin, *‘tsee!’*. In contrast, the spectrogram of the Southern Giant Hummingbird song had the shape of a candy cane (i.e., an inverted capital letter ‘J’; long and straight on one end, with a tight, curved bend at the top); by ear, it was recognizable as a loud ‘*tsiP!*’ with an abrupt ending.

### Song divergence between species

The Northern and Southern Giant Hummingbird differed strikingly in all measured song parameters. Importantly, song reliably identified the two cryptic species with consistently high confidence (Fig. 3). In PCA analyses of three bioacoustics traits (means of minimum frequency, maximum frequency, and song duration), PC1 corresponded strongly to species identity, explaining 80.9% variation in breeding data comparisons and 79.9% of the variation in full annual data comparisons (Fig. 3). PC2 explained variation in each minimum and maximum frequencies relative to song duration, explaining 11.5% variation in breeding data comparisons and 10.1% variation in full annual data comparisons.

**Figure 3.**
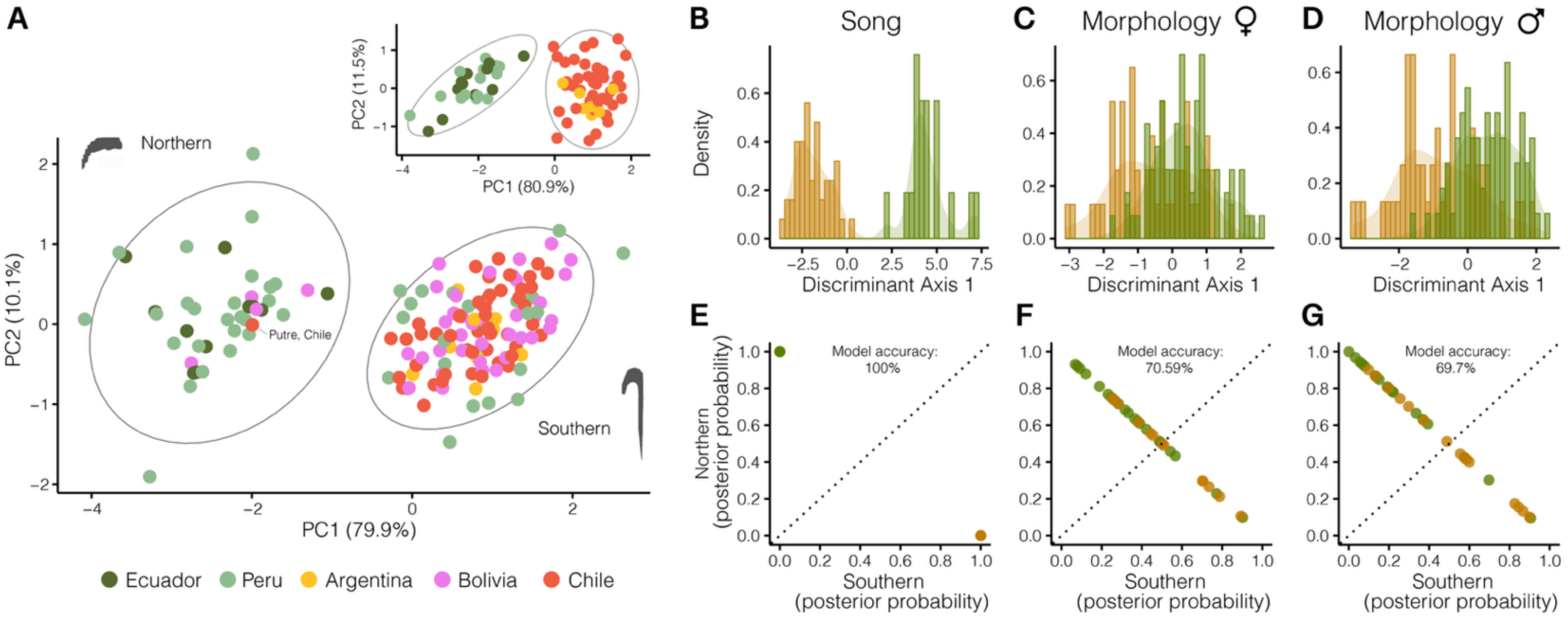
Divergent songs identify the cryptic giant hummingbird species. **A** Principal Component Analysis (PCA) of song characteristics (means of minimum frequency, maximum frequency, and duration) strongly separates species. Main plot shows full annual data, including non-breeding months when the two species co-occur (*n*=157 songs); inset shows breeding season only data (*n*=66 songs). **B** Linear Discriminant Analysis (LDA) of song characteristics (same as above; *n*=80 songs) identifies species with high confidence. **C,D** Results from LDA analyses of morphological characteristics (length of wing, tail, and bill; *n*=166 adult females and *n*=161 adult males), presented for comparison, highlight strong overlap in morphological characteristics. **E** The trained LDA bioacoustics model had 100% classification accuracy. **F–G** Trained LDA models for morphology of adult females and males classified species correctly in 70.59% and 69.7% of cases, respectively. Data in C–D and F–G are from Williamson et al. 2025b.

PCA results were consistent with the results of a LDA using minimum song frequency, maximum song frequency, and song duration. The trained LDA model predicted species identity with 100% classification accuracy using both training and test datasets (Fig. 3). However, one individual was misclassified (1.28% error rate) when the model was applied to full annual data. The individual in question was represented by ML631524705, a Southern Giant Hummingbird recorded by JLW on 12 February 2025 in Arani, Bolivia, that was classified by the model as a Northern Giant Hummingbird (Fig. S1). In all other cases, species identifications assigned by the trained LDA model matched those determined by assessment of season, locality, plumage, and molt pattern and timing.

### Song, more than morphology or plumage, reliably distinguishes species

In contrast to song, the distributions of Northern and Southern Giant Hummingbird morphological traits overlapped substantially (Fig. 3C–D). LDA models trained on previously published data of three morphological characteristics, and conducted separately for each adult females and males, could only distinguish Northern and Southern Giant Hummingbirds in 70.59% and 69.7% of instances, respectively (Williamson et al. 2024; Fig. 3 F–G). These results support previous findings that adult Northern and Southern Giant Hummingbirds may not always be reliably identified using morphological characteristics (Williamson et al. 2024, 2025) but highlight the relative ease with which song can distinguish the two.

### Clarified range limits and geographic co-occurrence

We recorded 38 individual giant hummingbirds of both species in Bolivia at 11 different localities, growing the existing collection of giant hummingbird songs archived in repositories by ∼75%. Our field recordings provided definitive evidence that the Northern Giant Hummingbird occurs in Bolivia; specifically, in the vicinity of Copacabana, Lake Titicaca (∼3,900 m), where we recorded four Northern Giant Hummingbirds (Fig. 1a). Three were adults and one was a juvenile; of the three adults, two were in fresh plumage and one bird was molting its outer primaries. No Northern Giant Hummingbirds were detected outside the vicinity of Copacabana, Bolivia, despite scouting in suitable *Patagona* habitat around other areas of La Paz Department, including Sorata, Pongo, the city of La Paz, Cañon de Palca, Bosquecillo Auquisamaña, and Achocalla, as well as throughout Cochabamba Department, where >62 Southern Giant Hummingbirds were observed in total.

We confirmed the occurrence of the Southern Giant Hummingbird in all sampled regions of Bolivia (Fig. 1a). Notably, we observed the Southern Giant Hummingbird around Copacabana, Bolivia, in the same sites where the Northern Giant Hummingbird was observed and recorded (at ∼3,900-4,000 m). All observed Southern Giant Hummingbirds were molting and were identifiable by their diagnostic molt patterns, combined with plumage and behavior. In some recordings, Northern and Southern Giant Hummingbirds were identified singing over one another.

## DISCUSSION

The Northern and Southern Giant Hummingbird differ markedly in their migration, genetics, and physiology, yet vary only subtly in external characteristics (Williamson et al. 2024, 2025). Until now, their vocal traits had not been formally compared. Our analyses revealed that the two species differ significantly in all measured song traits. Importantly, song appears to be the single most definitive diagnostic trait for identifying the two giant hummingbird species in the field. These findings are directly useful for ornithologists, birders, and naturalists observing and studying giant hummingbird ecology and evolution, as well as for management professionals conducting population-level censuses and conservation assessment. Because the song differences described here can be detected by ear (i.e., in the field or from recordings) and by eye (i.e., through visual assessment of spectrograms; Fig. 1), real-time or later inspection of spectrograms, provides an accessible way for hearing-impaired birders and researchers to identify hummingbird vocalizations.

That song can so easily distinguish the giant hummingbirds contrasts with the difficulty of identification with plumage or morphology (Fig. 3). Due to within-species trait variation and between-species overlap in external phenotypic characteristics, non-vocal individuals can be challenging to identify in the hand or in the field, even when scrutinized with the aid of trained statistical models, and even when considering data for each sex separately (Williamson et al. 2024, 2025). Whereas LDA models of morphological traits correctly identified adult females and males to species with 70.59% and 69.7% accuracy, respectively, LDA models trained on three song characteristics classified individuals correctly in 98.7–100% cases. Only one individual of 217 total was misidentified by the trained model when applied to full annual data (Fig. S1). This bird, represented by ML631524705, was included in the dataset despite having only one scorable song, partly because its identity was confirmed in the field by JLW; however, during scoring, RJZ noted that the recording for this individual appeared modulated. During field recording, JLW saw and heard this focal individual in an aerial chase with a second Southern Giant Hummingbird shortly after the single clear recorded song at ∼0:06 secs. These results highlight how field identification of the giant hummingbirds may require additional context clues, even with a diagnostic trait, such as song. Additionally, whereas morphological measurements and plumage may only be useful for distinguishing adult individuals (juvenile plumage of both species may be indistinguishable), song characteristics are applicable to both juveniles and adults.

The ranges of the giant hummingbirds have remained difficult to characterize using occurrence records and photos alone, and due to the previously undescribed migratory routes of some populations of the Southern Giant Hummingbird (Williamson et al. 2025). Our study provides new data on the ranges of both species in Bolivia and indicates a previously unknown zone of overlap. While preliminary given the limited scope of our field recording efforts, we nonetheless provide definitive evidence that the Northern Giant Hummingbird occurs in northern Bolivia. The southernmost extent of the Northern Giant Hummingbird range remains uncertain; however, we have not detected it south of Copacabana, Bolivia (Lake Titicaca region), along the eastern slope of the Andes. Interestingly, we documented that the Southern Giant Hummingbird occurs at the same high elevations and sites around Copacabana, Bolivia, consistent with previous genomic evidence (Williamson et al. 2024). At these sites, The Southern and Northern Giant Hummingbirds overlap, sometimes on the same hillside and within mere meters of each other. When recorded together, the two can be distinguished using audio and visual evidence (Figs. 1, 3). The Southern Giant Hummingbird thus appears to breed at high elevations in La Paz Department, a finding which awaits further substantiation. The northernmost extent of the Southern Giant Hummingbird breeding range remains unknown. Observers in the Lake Titicaca region of Peru and Bolivia can help expand our knowledge by recording songs of individuals observed during all months of the year. Follow-up research will be necessary to clarify breeding and non-breeding range limits and elevational limits, particularly during periods of the year when key floral resources (e.g., genera *Eucalyptus, Mutisia, Agave, Puya*) are in bloom.

Diagnostic songs present an opportunity to illuminate many aspects of giant hummingbird biology throughout the range of the genus. New field recordings from regions where few recordings exist (e.g., northwest Argentina, northern Chile) and from zones of sympatry will refine our understanding of seasonal ranges, elevations, and migratory phenology. For example, we identified several possible cases of apparent flexibility in the migration departure dates of the Southern Giant Hummingbird, relative to known tracked individuals from Chile: ML511071761 (recorded on 25 October 2022 in Cusco, Peru) may document a late- departing migratory Southern Giant Hummingbird, while XC17922 (recorded 9 March 2006 from Cusco, Peru) may document an early-arriving migratory Southern Giant Hummingbird; alternatively, these records may suggest that individuals originated outside of Chile, which remains to be confirmed.

In this study, we show that strikingly divergent songs identify the two giant hummingbird species with high confidence and provide a robust method for clarifying range limits and areas of seasonal co-occurrence. Our work additionally highlights the value of building and using open databases, such as Macaulay Library and Xeno-canto, in ecological and evolutionary research. Through the contributions of many recordists, we were able to analyze song data spanning 49 years, >36° of latitude, and >4,300 meters in elevation. The oldest recordings in our dataset were collected by ornithologist Ted Parker during a May 1976 trip to Ancash, Peru (Stap 1991). On this trip––which occurred during the giant hummingbird non-breeding season––Parker incidentally recorded both a Northern Giant Hummingbird (ML11057) and at least one Southern Giant Hummingbird (ML11062 and ML11063). That evidence of species-level divergence had remained hidden in the world’s largest repository of bird sounds dating back to 1976 evokes curiosity about potential cryptic species that have yet to be uncovered.

## Acknowledgements

We thank Eliot Miller and Vanessa Powell for assistance obtaining Macaulay Library recordings and metadata, Jay McGowan and Andrew Spencer for input on vocalizations, and Nicole Richardson and Carlos Carillo for assistance in the field. We are grateful to Jay McGowan for loaning equipment for song recording in the field. This work was supported by a Cornell Lab of Ornithology Ivy Fund Award to BWR, a Cornell Lab of Ornithology Experiential Learning Award to RJZ, and National Science Foundation (NSF-DBI-2208924) and Cornell Lab of Ornithology Edward W. Rose Postdoctoral Fellowships to JLW.

## Data Availability

Recordings are available in the public sound repositories Macaulay Library (https://www.macaulaylibrary.org/) Xeno-canto (www.xeno-canto.org). Data will be uploaded to Dryad upon manuscript acceptance.

