## Supplemental Figure S1 for "Songs distinguish the cryptic giant hummingbird species and clarify range limits"

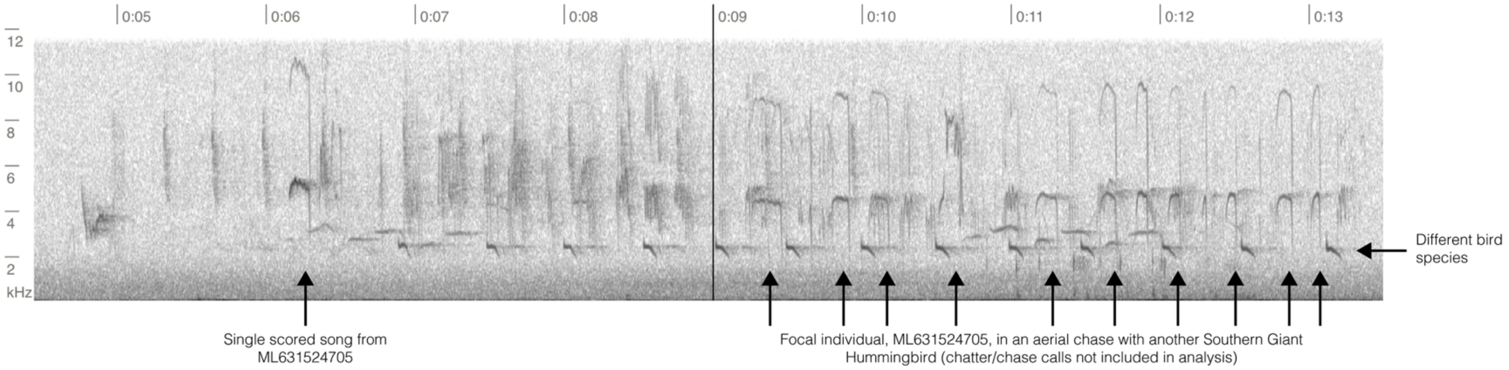


**Figure S1.** Excerpt of spectrogram recording from ML631524705 from 12 February 2025 in Cochabamba, Bolivia, the only individual that was misidentified by the linear discriminant model of vocal traits. The individual in question is a Southern Giant Hummingbird that was misidentified as a Northern Giant Hummingbird by the trained model. Only one song was scored for this individual; during scoring, RJZ noted that the song appeared modulated (note "wavery" top loop of typical Southern candy cane shape). In the rest of the recording, this individual can be heard fighting with second Southern Giant Hummingbird in an aerial chase. JLW observed the focal individual, as well as the individual it fought, for an extended period and additionally took diagnostic plumage and molt photos. All data support the ID of Southern Giant Hummingbird. The ML record is available at: https://ebird.org/checklist/S213020346.
